# Coordinated Temporal Dynamics of Glucocorticoid Receptor Binding and Chromatin Landscape Drive Transcriptional Regulation

**DOI:** 10.64898/2026.04.23.720290

**Authors:** Diana A. Stavreva, Sohyoung Kim, Saori Fujiwara, Andrew McGowan, Songjoon Baek, Lorenzo Rinaldi, Thomas A. Johnson, Michele Puglia, Blagoy Blagoev, Franck Dequiedt, Gordon L. Hager, Gregory Fettweis

## Abstract

Glucocorticoid receptor (GR) signaling elicits diverse transcriptional responses through dynamic and context-dependent interactions with chromatin. Here, we define a temporally resolved and mechanistically integrated framework for GR-mediated gene regulation. Time-resolved analyses identify three conserved classes of GR chromatin binding (sustained, transient, and late), distinguished by differences in motif strength, chromatin accessibility, and cofactors engagement. Early GR binding preferentially occurs at high-affinity glucocorticoid response elements (GREs) within pre-accessible regulatory regions, whereas late binding is associated with weaker motifs and requires chromatin remodeling activity. Enhancer activation, marked by H3K27ac deposition, closely tracks GR occupancy, supporting a model in which GR recruits acetyltransferase activity to drive coordinated enhancer activation. Concurrently, GR-centered interaction networks are dynamically reconfigured, and motif enrichment analyses identify distinct transcription factor signatures across binding classes, including AP-1/JUNB at transient sites and CEBP family members at late-binding regions. Integration of chromatin binding, chromatin interaction, and transcriptomic datasets reveals that temporal and combinatorial GR occupancy is functionally linked to gene expression programs. Distinct GR binding clusters are nonrandomly associated with specific transcriptional trajectories, including sustained, transient, and late gene induction. Moreover, combinatorial occupancy across multiple regulatory elements correlates quantitatively with transcriptional output, indicating that GR functions not as a simple binary regulator, but as an integrator of multilayered regulatory inputs.

These findings support a unified model in which temporal binding dynamics, chromatin state, and combinatorial enhancer activity collectively encode transcriptional specificity, providing a general framework for stimulus-responsive nuclear receptor signaling.

## Introduction

Glucocorticoids regulate diverse physiological processes, including inflammation, metabolism, development, and stress responses, through signaling mediated by the glucocorticoid receptor (GR), a ligand-activated transcription factor of the nuclear receptor superfamily. Upon ligand binding, GR translocates to the nucleus, where it coordinates transcriptional programs that enable cells and tissues to respond to hormonal and environmental stress signals. GR modulates gene expression by binding to glucocorticoid response elements (GREs) and assembling transcriptional regulatory complexes with coactivators, corepressors, and chromatin-remodeling factors^1,2^.

GR-mediated transcription is highly complex and context-dependent. Time-resolved transcriptomic analyses reveal that most GR-responsive genes exhibit dynamic, multiphasic expression patterns rather than simple monotonic activation or repression. GR-regulated genes often display transient induction, delayed activation, or alternating phases of induction and repression. Thus, glucocorticoid signaling unfolds through temporally coordinated regulatory programs rather than binary gene regulation^3^. Whereas large-scale analyses have shown that GR binds to thousands of genomic sites, only a subset of these binding events is associated with measurable transcriptional responses. This observation indicates that additional regulatory mechanisms determine the transcriptional outcome of GR binding^4,5^.

Chromatin organization plays a central role in shaping GR signaling across the genome. The chromatin landscape strongly constrains GR occupancy and contributes to cell type-specific glucocorticoid responses^6^. Consistent with this model, lineage-determining transcription factors can function as chromatin-priming factors that facilitate GR recruitment to specific regulatory elements^7^. GR binding loci often coincide with distal regulatory elements functioning as enhancers rather than promoters^6,7^. GR-mediated activation of these enhancers is associated with the recruitment of transcriptional coactivators such as p300 and CBP, which catalyze histone H3 lysine 27 acetylation (H3K27ac) and promote transcriptional activation^8,9^.

At the molecular level, GR operates within highly dynamic transcriptional complexes. Live-cell imaging studies have revealed rapid GR exchange at chromatin-binding sites^10^, while hormone stimulation modulates transcriptional bursting at GR target loci^11^. These dynamic interactions, together with cyclical recruitment of transcriptional cofactors and RNA polymerase II, contribute to the complex temporal patterns of gene expression observed following hormone stimulation^3,12^.

Mechanistic insights into these diverse transcriptional responses are particularly important because glucocorticoids remain among the most widely prescribed anti-inflammatory and immunosuppressive drugs in clinical medicine^13,14^. However, their therapeutic benefits are frequently accompanied by significant adverse effects and variable responsiveness across tissues and disease contexts^15,16^. Defining how chromatin architecture, transcription factor networks, and enhancer dynamics shape GR-mediated transcriptional programs will therefore be essential for improving the specificity and therapeutic index of glucocorticoid-based treatments.

## Materials and Methods

### Cell culture and treatment

C127 cells (ATCC CRL-1616) or the derivative 33AP1C9 cell line used in the Pol II-ChIP experiments^11^ were cultured in Dulbecco’s modified Eagle’s medium (DMEM) supplemented with 10% fetal bovine serum (FBS), sodium pyruvate, L-glutamine, and nonessential amino acids, and maintained at 37 °C in 5% CO_2_.

For hormone treatment experiments, cells were cultured in DMEM supplemented with 5% charcoal-stripped FBS for at least 48 h prior to treatment. Cells were then treated with 600 nM corticosterone for either 1 h or 12 h. Control cells were treated with ethanol (vehicle; Eth.) for 1 h.

### Antibodies

Primary antibodies were as follows: anti-GR (Santa Cruz Biotechnology sc-393232), anti-H3K27ac (Abcam, ab4729), anti-H3K4me1 (Abcam, ab8895), anti-RPB1 [RNA polymerase II CTD phospho S5 (Abcam, ab5131)].

### ChIP-seq and ATAC-seq

ChIP-seq and ATAC-seq were performed as previously described^17^. C127 cells were plated on 15 cm dishes (3 million cells per dish) and grown for 72 h in stripped medium. Cells were treated with vehicle (EtOH) or 600 nM corticosterone for 1 h or 12 h prior to ChIP or nuclei isolation for ATAC-seq.

For chromatin immunoprecipitation (ChIP), cells were crosslinked with paraformaldehyde and subsequently harvested. Chromatin was sonicated using a Bioruptor system (Diagenode) to obtain an average DNA fragment size of 200-500 bp. For immunoprecipitation, 800 µg of chromatin was incubated overnight at 4 °C with the appropriate antibody pre-bound to Dynabeads magnetic beads (Thermo Fisher Scientific) under continuous rotation. Following stringent washes, antibody-bound chromatin complexes were eluted, cross-links were reversed, and proteins were enzymatically digested. DNA was purified by phenol-chloroform extraction followed by ethanol precipitation. ChIP-seq libraries were prepared using the TruSeq ChIP Sample Preparation Kit (Illumina, IP-202-1012) according to the manufacturer’s protocol. Antibodies used for immunoprecipitations: GR, 4 µg; H3K27ac, 2 µg; H3K4me1, 2 µg; RPB1, 2 µg.

ATAC-seq was performed following the Omni-ATAC protocol^18,19^. Briefly, 50,000 viable fresh cells were collected and resuspended in ATAC resuspension buffer (10 mM Tris-HCl [pH 7.4], 10 mM NaCl, 3 mM MgCl_2_) supplemented with 0.1% NP40, 0.1% Tween-20, and 0.01% digitonin. The cell suspension was incubated on ice for 3 minutes. Subsequently, a wash step was carried out using ATAC resuspension buffer containing 0.1% Tween-20, followed by centrifugation at 4°C for 10 minutes. After removal of the supernatant, isolated nuclei were resuspended in Omni-ATAC transposition mix containing Omni-TD buffer (10 mM Tris-HCl [pH 7.6], 5 mM MgCl_2_, 10% dimethylformamide), TDE1 transposase (Illumina, 20034197), 0.1% Tween-20, 0.01% digitonin, and 1× PBS, then incubated at 37°C for 30 minutes with agitation (1000 rpm) in a thermomixer. Tagmented DNA was purified using the MinElute PCR Purification Kit (Qiagen, 28004). Samples were then amplified by PCR using NEBNext High-Fidelity PCR Master Mix (New England Biolabs, M0541S) and custom Nextera indexing primers^20^, with a total of 8-9 amplification cycles. Amplified libraries were purified and size-selected (150-1000 bp) using SPRIselect beads (Beckman Coulter, B23317), and DNA was eluted in 10 mM Tris-HCl (pH 8.5). Library quality and fragment size distribution were assessed using D1000 and D5000 ScreenTape assays (Agilent Technologies). Finally, pooled libraries were sequenced on an Illumina NextSeq platform using paired-end reads.

### Next-generation sequencing data Analysis

ChIP-seq and ATAC-seq read quality filtering and alignment to mouse (mm10) or human (hg38) genomes were performed as previously described. Downstream analyses were conducted using HOMER^21^ as described previously^17^.

For experiments in this study, GR peak calling was performed using the *findPeaks*.*pl* script with the following parameters: FDR < 0.00001, fold change compared to control = 6, local fold change = 6, and tagThreshold = 100. For the d’Ippolito dataset, GR peak calling was performed with the same parameters, except tagThreshold = 50. GR peak clusters (C1/DC1, C2/DC2, and C3/DC3) were defined using the *mergePeaks*.*pl* script. Heatmaps and aggregate plots were generated using 20 bp bins across ±1-2 kb regions centered on peak summits. All plots were normalized to 10 million mapped reads and further normalized to local tag density (tags per base pair per site). *De novo* motif analysis was performed using *findMotifsGenome*.*pl*. Box plots were generated, and statistical significance was assessed using one-way ANOVA with a Bonferroni post hoc test.

### Determination of GR ChIP-Seq peaks linked to GR-Dependent genes

A total of 638 genes were identified using RefSeq IDs and gene symbols. Gene boundaries were defined as gene bodies extended by 2 kb upstream of the transcription start site, based on integrated annotations from the HOMER database (v.5.1; mm10.tss, mm10.rna, mm10.basic.annotation and mouse2gene.tsv). GR peaks located within 50 kb of these gene boundaries were assigned as putative regulatory elements. In addition, distal GR peaks located at loop anchors were assigned to genes if one anchor overlapped a GR peak and the interacting anchor was located within 50 kb of the gene boundary, based on chromatin interactions identified by GR HiChIP^22^. Interactions involving up to two sequential looping events were included. In total, 1,616 gene-GR peak associations were identified, involving 482 genes and 1,128 GR peaks.

### Correlation analysis between GR chromatin binding and GR transcripts profile

Based on gene-GR peak assignments, we summarized GR peak patterns for each gene using three peak modes: C1 (1 h, 12 h), C2 (1 h), and C3 (12 h). For each gene, a relative frequency vector (C1, C2, C3) was calculated across all associated GR peaks. For example, A (0.25, 0.75, 0) indicates that 25% and 75% of GR peaks assigned to gene A belong to the C1 and C2 modes, respectively.

Each gene frequency vector(n = 482) were subjected to hierarchical clustering using pheatmap R package (v1.0.12) with default parameters, resulting in five major pattern classes comprising 187, 184, 76, 18, and 13 genes. To assess the association between the six gene expression patterns (sustained, transient and delayed induction; sustained, transient and delayed repression) and these GR peak pattern classes, we performed Pearson’s chi-squared tests for each peak pattern class, followed by post hoc analysis using chisq.posthoc.test R package (v0.1.3). Results are reported as FDR-adjusted P-values (Benjamini-Hochberg) and Pearson residuals. All analyses were performed in R (v4.3.2).

### Bivariate Analysis of Genomic Footprint (BAGFoot)

Genome-wide differential chromatin accessibility and footprinting were analyzed using Bivariate Analysis of Genomic Footprinting^23^ (BaGFoot; R version 0.9.7) with default parameters to identify transcription factor (TF) motifs showing occupancy changes across conditions.

ATAC-seq peaks across the three conditions (Eth., 1 h, and 12 h) were used to quantify cut-site frequencies. For each TF motif, accessibility and footprint depth scores were computed, and pairwise comparisons were performed to derive differential accessibility and footprint depth.

### RNA extraction, sequencing and analysis

C127 cells were treated with 600 nM corticosterone for 1 h or 12 h prior to RNA isolation. RNA was isolated using the PureLink RNA Kit (Thermo Fisher Scientific, 12183018A) according to the manufacturer’s instructions. RNA-seq libraries were generated from rRNA-depleted total RNA samples (Illumina, RS-122-2301) using the Illumina TruSeq Stranded Total RNA Library Prep kit (Illumina, 20020596), following the manufacturer’s protocol. Libraries were sequenced on the HiSeq3000/4000 platform to generate 2 x 150 bp paired-end reads. Base calling was performed using Illumina RTA (v1.18.64), and reads were demultiplexed using Illumina bcl2fastq (v2.17) to generate sample specific FASTQ files. Samples yielded 69-85 million pass-filter reads, with >91% of bases achieving a quality score ≥ Q30. Sequencing reads were trimmed for adaptor sequences and low-quality bases using Trimmomatic (v0.36). Trimmed reads were aligned to the mouse reference genome (mm10) with GENCODE vM9 annotations using STAR (v2.5.1) in two-pass mode. Three biological replicates were sequenced for each condition. Downstream analysis was conducted using the HOMER intronic pipeline^21^ and DESeq2^24^. Differentially expressed genes were considered significant with a fold change ≥ 2 and an FDR corrected p-value ≤ 0.05.

### GR *ChIP-SICAP proteomic*

All proteomic experimental procedures and peptide discovery has already been described in detail in our previous work^22^. In brief, A spectral library was generated from 20% of trypsin-digested ChIP-SICAP peptides using high-pH fractionation and DDA LC-MS/MS on a Q Exactive HF-X system. Raw data were processed in MaxQuant with standard parameters (trypsin digestion, common modifications, 1% FDR) against a *Mus musculus* database. The remaining 80% of peptides were analyzed by DIA using the generated spectral library, as described^25^. DIA data were processed in Skyline with strict mass accuracy, retention time filtering, and mProphet scoring (Q < 0.01). Quantified data were normalized, and differential interactions were defined based on ≥ 50% sample presence and ≥ 2-fold change across GR conditions.

### Data availability

RNA-seq, ChIP-seq data, and ATAC-seq generated for this study were deposited to the NCBI Gene Expression Omnibus (GEO). The ENCODE data from the Reddy laboratory^26^ used in this study are gathered here (https://www.encodeproject.org/awards/U01HG007900/). The TF motifs used in this manuscript can be found at the HOMER database (http://homer.ucsd.edu/homer/). All other relevant data are available from the corresponding author upon request.

## Results

### Corticosterone treatment reveals distinct temporal classes of GR-chromatin binding

To characterize the dynamics of glucocorticoid receptor (GR) chromatin binding, we performed GR ChIP-seq at baseline (ethanol-treated cells, Eth.), 1 h, and 12 h, following corticosterone (Cort) treatment. Clustering analysis revealed three major classes of GR peaks based on temporal occupancy patterns: C1 (sustained binding), occupied at both 1 h and 12 h; C2 (transient binding), bound exclusively at 1 h; and C3 (late-binding), observed only at 12 h (Fig. 1A). Quantification of GR occupancy using HOMER tag density confirmed that C1 sites exhibit the strongest and most persistent GR binding across both time points, whereas C2 sites display robust but transient enrichment at 1 h, and C3 sites show comparatively weaker binding emerging at 12 h (Fig. 1B). Aggregate GR ChIP-seq profiles confirmed these patterns, with C1 maintaining high signal intensity over time, while C2 and C3 exhibited time-restricted binding profiles (Fig. 1C).

**Figure 1.**
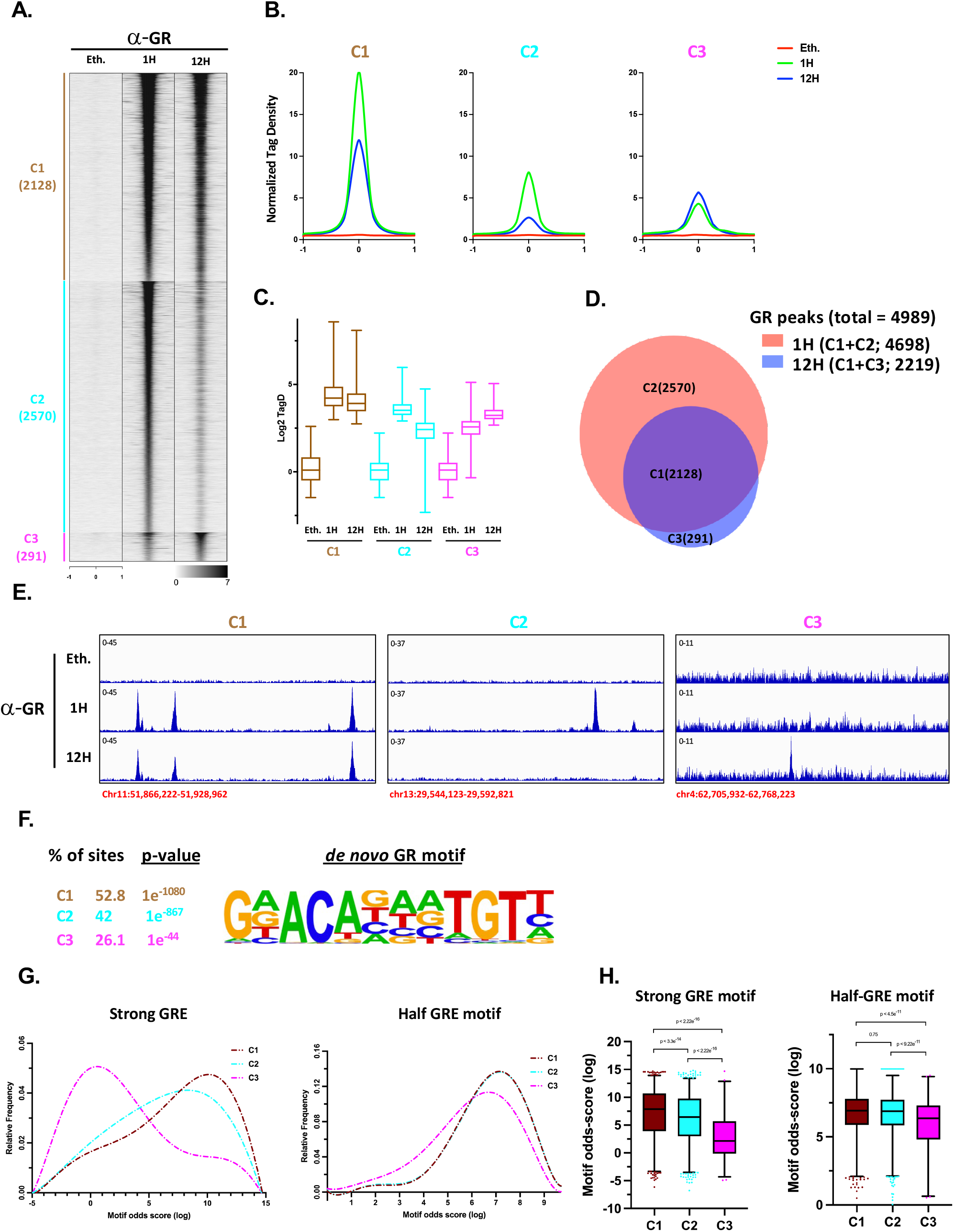
Temporal classes of GR chromatin binding following glucocorticoid receptor activation. **(A)** GR ChIP-seq signal heatmap in C127 cells treated with Ethanol (Eth.) and with corticosterone 600 nM for 1 h, and 12 h. Peaks are grouped into three clusters according to peak calling parameters (see method): C1, detection at 1 h and 12 h; C2, detection only at 1 h; and C3, detection only at 12 h. Peak numbers per cluster are indicated. Data are displayed ±1 kb from peaks center. Binding intensity (tags per base pair per site) scale is noted below on a linear scale. (**B**) Aggregate plots of GR ChIP-seq cluster profiles (20-bp bins) around peak center. (**C**) Normalized log_2_ TagDensity across GR clusters. (**D**) Venn diagram of GR ChIP-seq clusters. (**E**) Representative genome browser tracks illustrating GR binding patterns for C1, C2, and C3 clusters. (**F**) Top enrichment GR motif from a *de novo* motif analysis of all GR peaks (n = 4,989). The enrichment is displayed as percentage of the number of sites with the calculated p-values. (**G**) Analysis of canonical (strong) and half-site (weak) GR response elements motifs odds scores (log scale) and frequencies were calculated for each GR cluster. (**H**) Boxplots of motif odds-score distributions from panel G (KW-test) p-val<0.05 is considered significant. See also Supplementary Figure 1

Approximately twofold more GR peaks were detected at 1 h (n=4698) compared to 12 h (n=2219) (Fig. 1D). Comparison of peaks between time points revealed a large overlap, indicating that most GR binding events occur early after hormone treatment. Representative genome browser tracks further illustrate the distinct temporal binding behaviors of each cluster (Fig. 1E).

Genomic annotation of GR binding sites revealed that peaks from all clusters are predominantly located in intergenic and intronic regions, consistent with distal regulatory elements (Fig. S1A). No major differences in genomic distribution were observed among the clusters. Analysis of peak proximity indicated that GR binding sites frequently occur in clusters across the genome. A substantial fraction of peaks has at least one neighboring GR site, defined as the presence of another peak within ±50 kb. This pattern is consistent across clusters (Fig. S1B-C). However, distance-based analyses revealed subtle differences : C3 peaks tend to be located closer to neighboring GR sites compared to C1 and C2, as reflected in cumulative distribution and nearest-neighbor analyses (Fig. S1D-E). Overall distributions of inter-peak distances and the number of neighboring peaks per site are comparable across clusters (Fig. S1F-G). Overall, GR binding sites exhibit similar genomic distribution and clustering patterns across all groups, with only minor differences in local spacing.

*De novo* motif analysis (n = 4,989) identified the canonical glucocorticoid response element (GRE) as the most significantly enriched motif in GR peaks (Fig. 1F). Cluster-specific analysis revealed strong GRE enrichment in C1 and C2 sites, whereas C3 sites showed comparatively weaker enrichment. To further assess GR motif quality, we quantified the presence of canonical (strong) GREs and half-site (weak) GREs across clusters. Motif odds scores and frequencies indicated that C1 sites are the most enriched for high-affinity canonical GREs, followed by C2 and then C3 (Fig. 1G).

Statistical analysis using Kruskal-Wallis tests confirmed significant differences in canonical GRE enrichment across all clusters, with C1 showing the highest motif strength (Fig. 1H). In contrast, enrichment in weak GREs was similar in C1 and C2 and only modestly reduced in C3.

To assess reproducibility, we compared our dataset with previously published GR ChIP-seq data from D’Ippolito et al.^26^. Applying our analytical framework, GR peaks were classified into distinct groups based on temporal binding dynamics: DC1 (sustained), DC2 (transient), and DC3 (late-binding). Consistent with our findings, we uncovered a similar triad of GR chromatin binding patterns in the human A549 cells with the transient C2 cluster (DC2 for this dataset) representing the largest fraction of GR binding events (Fig. S1H). Similar temporal overlap patterns between 1 h and 12 h peaks were also observed (Fig. S1I), as well as comparable motif enrichment trends, with strong GRE enrichment in early/transient sites and weaker enrichment in late-binding sites (Fig. S1J). Together, these findings reveal a conserved temporal architecture of GR chromatin binding across species and cell lines and identify the GR binding motif as a key determinant of this pattern.

### Specific chromatin contexts underlie GR temporal chromatin binding patterns

To investigate how chromatin context relates to temporal patterns of GR binding, we profiled chromatin accessibility using ATAC-seq and histone modifications associated with active enhancers (H3K27Ac and H3K4me1) at GR binding sites across the distinct temporal clusters. Heatmap visualization of ATAC-seq, H3K27ac, H3K4me1, and RNA polymerase II (Pol II) ChIP-seq signals revealed specific chromatin features associated with each cluster (Fig. 2A). Indeed, the early binding sites (C1 and C2) exhibited strong pre-existing chromatin accessibility and active histone marks, whereas late-binding sites (C3) showed comparatively stronger signals prior to stimulation, suggesting a more open chromatin state before GR recruitment.

**Figure 2.**
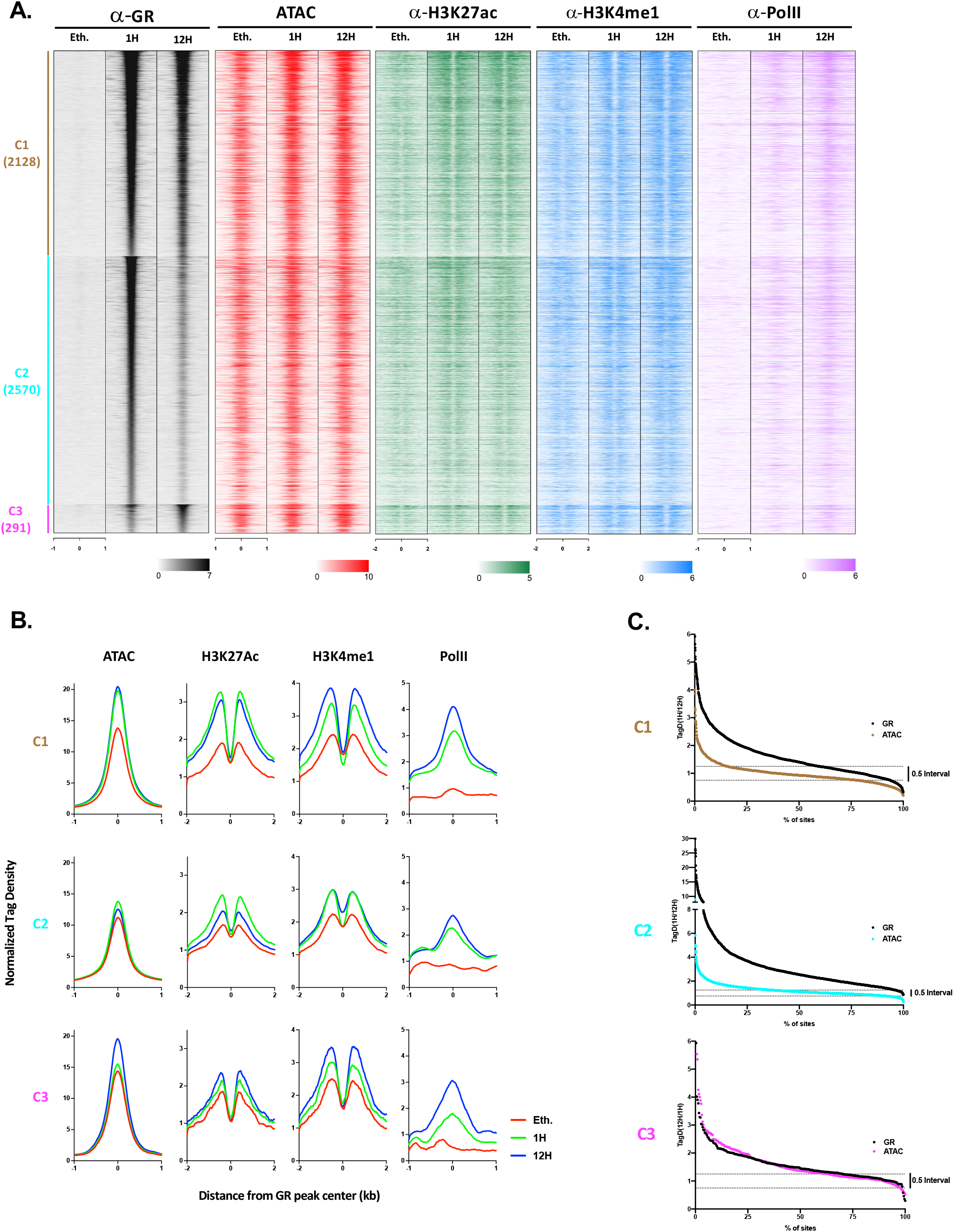
Chromatin landscape associated with temporal GR binding classes. (**A**) Heatmaps of ATAC-seq, H3K27ac, H3K4me1, and Pol II Chip-seq signals at GR binding sites, shown ±1 or 2 kb from peaks center and organized by GR temporal clusters (C1-C3). (**B**) Aggregate plots related to panel 2A for each cluster profiles (20-bp bins) around peaks center. The Kruskal–Wallis test was used to evaluate differences among treatment conditions; only statistically significant results are reported (pval<0.05). (**C**) Comparison of ATAC-seq and GR ChIP-seq TagDensity dynamics between 1 h and 12 h Cort treatment. Time related ratios for each GR peak are shown as cumulative distribution functions (CDFs). See also Supplementary Figure 2

Aggregate signal profiles confirmed these trends and highlighted dynamic changes following corticosterone (Cort) treatment (Fig. 2B). In a distinct manner, each cluster displayed a specific dynamic pattern of chromatin features. In C1, chromatin accessibility was significantly increased relative to baseline; however, no significant difference was observed between 1 h and 12 h of treatment. A similar pattern was also visible for the H3K27ac mark. In contrast, H3K4me1 exhibited a more gradual increase from the ethanol condition to 12 h of treatment. Conversely, in the C2 cluster, chromatin accessibility did not change upon treatment, while H3K27ac marks showed a significant but transient increase at 1 h of treatment. H3K4me1 levels were only weakly induced by the treatment and remained relatively stable across the 1 h and 12 h time points. In C3, chromatin accessibility remained unchanged after treatment, whereas both H3K27ac and H3K4me1 marks displayed a gradual increase over time.

In contrast to the dynamic changes observed for H3K27ac, RNA polymerase II (Pol II Ser5P) occupancy displayed a progressive increase over time from 0 h to 12 h across all GR binding clusters (Fig. 2A, B).

This accumulation occurred irrespective of the distinct temporal patterns of GR binding and H3K27ac enrichment. In particular, while H3K27ac closely paralleled GR occupancy, showing sustained enrichment in C1, transient induction in C2, and delayed accumulation in C3, Pol II signals increased continuously even in clusters where H3K27ac plateaued or declined at later time points.

Focusing specifically on intergenic GR binding sites, heatmaps and aggregate profiles of Pol II ChIP-seq signal across intergenic clusters (iC1–iC3) recapitulated the patterns observed across all GR peaks (Supplementary Fig. 2C, D), indicating that Pol II enrichment is not solely driven by proximity to promoters but reflects a broader transcriptional engagement at distal regulatory elements. In agreement, eRNA production at intergenic sites followed the temporal pattern of Pol II accumulation (Supplementary Fig. 2D). Notably, this progressive behavior parallels the gradual increase observed for H3K4me1, suggesting that transcriptional engagement and enhancer maturation are cumulative processes, in contrast to the transient nature of H3K27ac.

Thus, among the chromatin features examined, H3K27ac most closely parallels the temporal GR binding patterns defined previously, showing rapid enrichment at early GR-binding sites and delayed accumulation at late binding regions. This suggests that enhancer activity, as marked by H3K27ac, is tightly correlated to GR occupancy.

Consistent with these findings, profiling chromatin features across GR temporal clusters in the D’Ippolito dataset revealed distinct regulatory signatures, including differential ATAC-seq signals and H3K27ac enrichment at enhancer-associated regions (Supplementary Fig. 2A-B). Notably, EP300, the histone acetyltransferase responsible for H3K27 acetylation, exhibited dynamics that closely mirror those of H3K27ac, further supporting a tight coupling between H3K27ac deposition and GR chromatin binding dynamics.

To better characterize the relationship between chromatin accessibility and GR binding, we further analyzed changes in ATAC-seq and GR ChIP-seq signals between early (1 h) and late (12 h) time points. Cumulative distribution analyses showed that C1 and C2 sites underwent relatively modest changes in accessibility despite substantial differences in GR occupancy (Fig. 2C), indicating that GR binding at these sites occurs largely within pre-accessible chromatin. In contrast, C3 sites displayed coordinated increases in both ATAC-seq and GR signal, consistent with a model in which chromatin remodeling facilitates GR recruitment at late-binding loci.

These results support a model in which temporal GR binding patterns correspond to distinct chromatin states: early-binding sites are located within pre-accessible, active enhancers, whereas late-binding sites require chromatin remodeling, with H3K27ac dynamics most closely tracking GR occupancy and enhancer activation.

### GR protein interaction network is dynamically remodeled over time

To investigate temporal changes in GR protein interactions, we performed GR ChIP-SICAP proteomic analyses^27^ following corticosterone treatment (Fig. 3A). Comparative analysis revealed a substantial reduction in the number of GR-associated proteins at 12 h relative to 1 h, suggesting that GR chromatin-associated complexes undergo dynamic remodeling over time after an initial increase of protein-protein interactions (Fig. 3B). Quantitative analysis of protein enrichment relative to the 1 h condition identified distinct patterns of interaction dynamics (Fig. 3C). Clustering of these interaction-profiles revealed three major groups: sustained interactors that remained associated over time, transient interactors that peaked at 1 h and declined by 12 h, and a small subset of late interactors that appeared only at the later time point (Fig. 3D).

**Figure 3.**
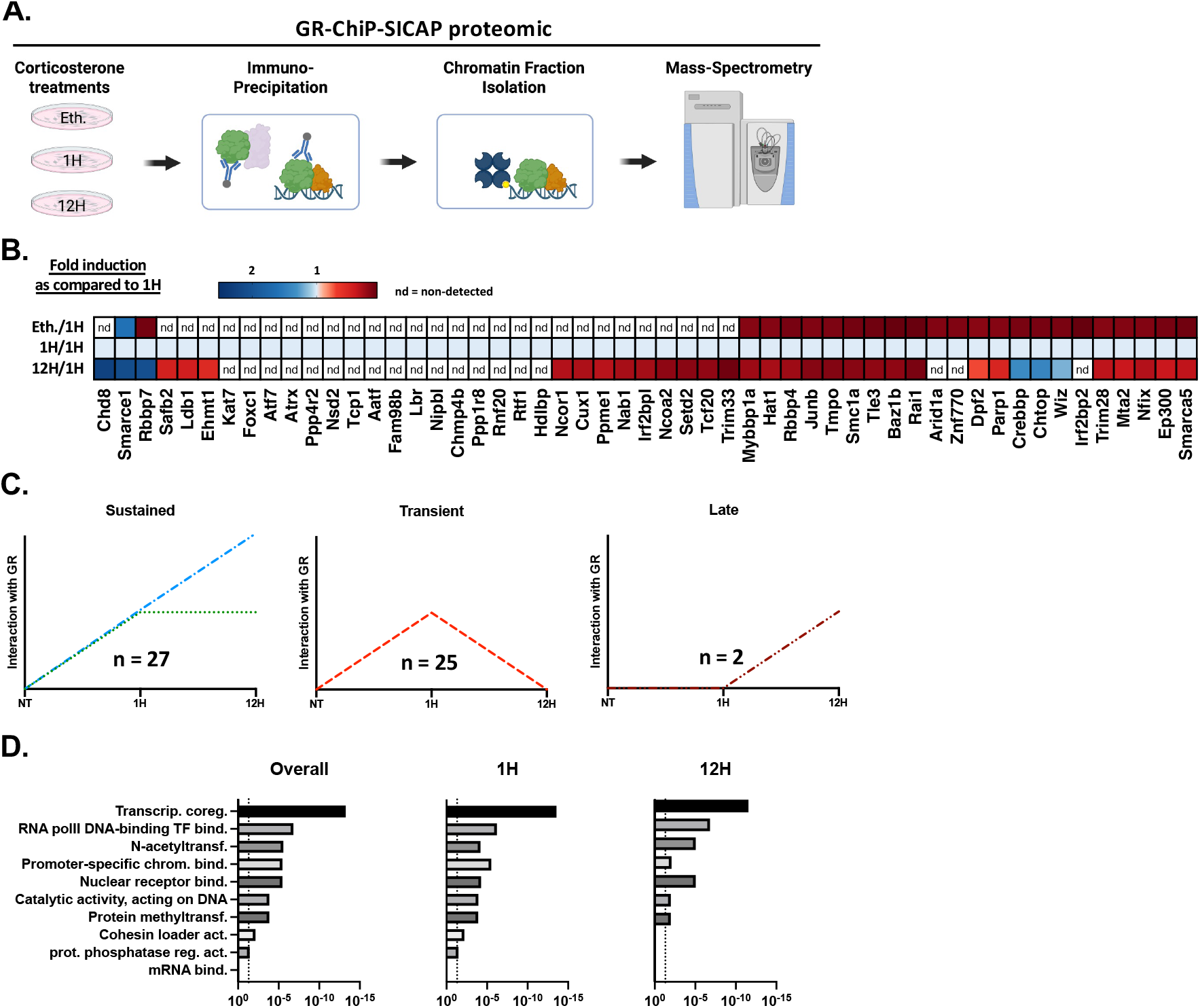
GR protein interaction network is time-dependent. (**A**) Experimental strategy for GR Chip-SICAP proteomic experiments. (**B**) Heatmap of abundance ratio of peptides detected after 1 h of corticosterone as compared to ethanol (Eth.) and 12 h of treatments. (**C**) GR interactants temporal association pattern. (**D**) Molecular function gene ontology enrichment of GR interactants. The dash line represents a p-value of 0.05.

Gene ontology (GO) enrichment analysis of GR interactants demonstrated significant overrepresentation of transcription-related functions, including “transcriptional co-regulator activity” (indicate the GOs), “RNA polymerase II transcription factor binding” (indicate the GOs), and chromatin-associated enzymatic processes (indicate the GOs) (p < 0.05; Fig. 3E). Notably, early time points were enriched for a broader range of transcriptional regulators, whereas later time points showed a more restricted functional profile whith the loss of somes significant GO classifications.

Together, these findings indicate that GR chromatin-associated protein interactions are dynamically remodeled over time, with an overall reduction after the initial response and reveal a complex, evolving interactome comprising multiple transcription factors, suggesting distinct TF co-binding mechanisms across different GR peaks.

### Temporal GR binding clusters exhibit distinct transcription factor motif enrichment

To identify transcription factor networks associated with temporal GR binding, we performed *de novo* motif discovery separately for each GR temporal cluster using HOMER. Analysis was conducted on sequences centered on GR peaks, allowing identification of cluster-specific regulatory signatures. Distinct non-GR transcription factor motifs were enriched across clusters, with the top motifs in each cluster represented by the percentage of peaks containing the motif and their associated statistical significance (Fig. 4A). Notably, ATF3, JUNB, and CEBPα motifs emerged as the most significantly enriched motifs across clusters. JUNB motifs were preferentially enriched in transient GR binding sites (C2), whereas CEBPα motifs were selectively enriched in late-binding sites (C3), indicating differential cofactor involvement depending on GR binding dynamics. In contrast, ATF3 motifs were broadly enriched, with a bias toward early or sustained binding sites (C1), suggesting a potential role in initial GR recruitment or early transcriptional responses.

**Figure 4.**
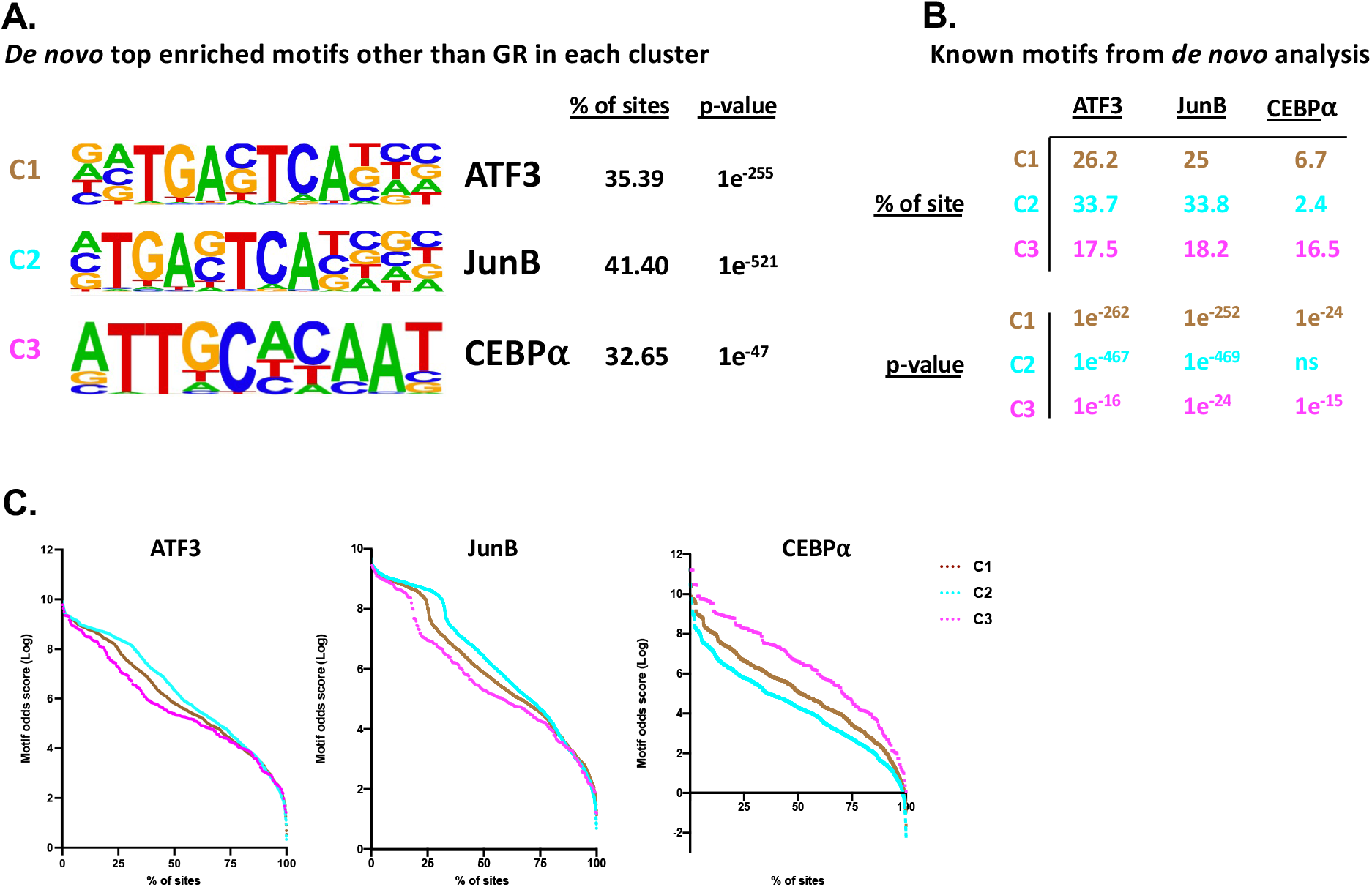
Cluster-specific transcription factor motif enrichment at GR binding sites. (**A**) Top enriched non-GR motifs identified in each cluster by *de novo* motif analysis, shown as the percentage of peaks containing each motif with associated p-values. (**B**) Enrichment of the motifs shown in panel 4A across the GR clusters, displayed as percentage of peaks and p-values. (**C**) Cumulative distribution functions (CDFs) of motif odds scores for each motif across GR peaks within each cluster.

Comparison of motif enrichment across all clusters confirmed the specificity of these associations, as motifs identified in individual clusters displayed distinct enrichment patterns when evaluated across all GR binding sites (Fig. 4B). JUNB and ATF3 motifs showed strong enrichment in early and transient clusters, whereas CEBPα motifs exhibited comparatively higher enrichment in late-binding sites, reinforcing the temporal specificity of these transcription factor associations.

Consistent with these findings, cumulative distribution function (CDF) analysis of motif odds scores revealed clear shifts in motif score distributions across clusters (Fig. 4C). JUNB and ATF3 motifs showed higher motif scores in early and transient clusters, while CEBPα motifs displayed increased scores in late-binding sites. Together, these results demonstrate that GR binding dynamics are associated with distinct transcription factor motif landscapes, supporting a model in which GR cooperates with different cofactors over time to regulate stage-specific gene expression programs.

### Coordinated changes in chromatin accessibility and transcription factor occupancy

To further validate our findings, we performed *de novo* motif analysis of GR temporal clusters in the previously published D’Ippolito dataset^26^. Consistent with the motif analysis results from our dataset (Fig. 4) and previously published data^28,17,29^, the analysis confirmed enrichment of key non-GRE transcription factor binding sites, including AP-1 family members (c-Jun/JunB) and CEBPα, across corresponding clusters (Fig. 5A).

**Figure 5.**
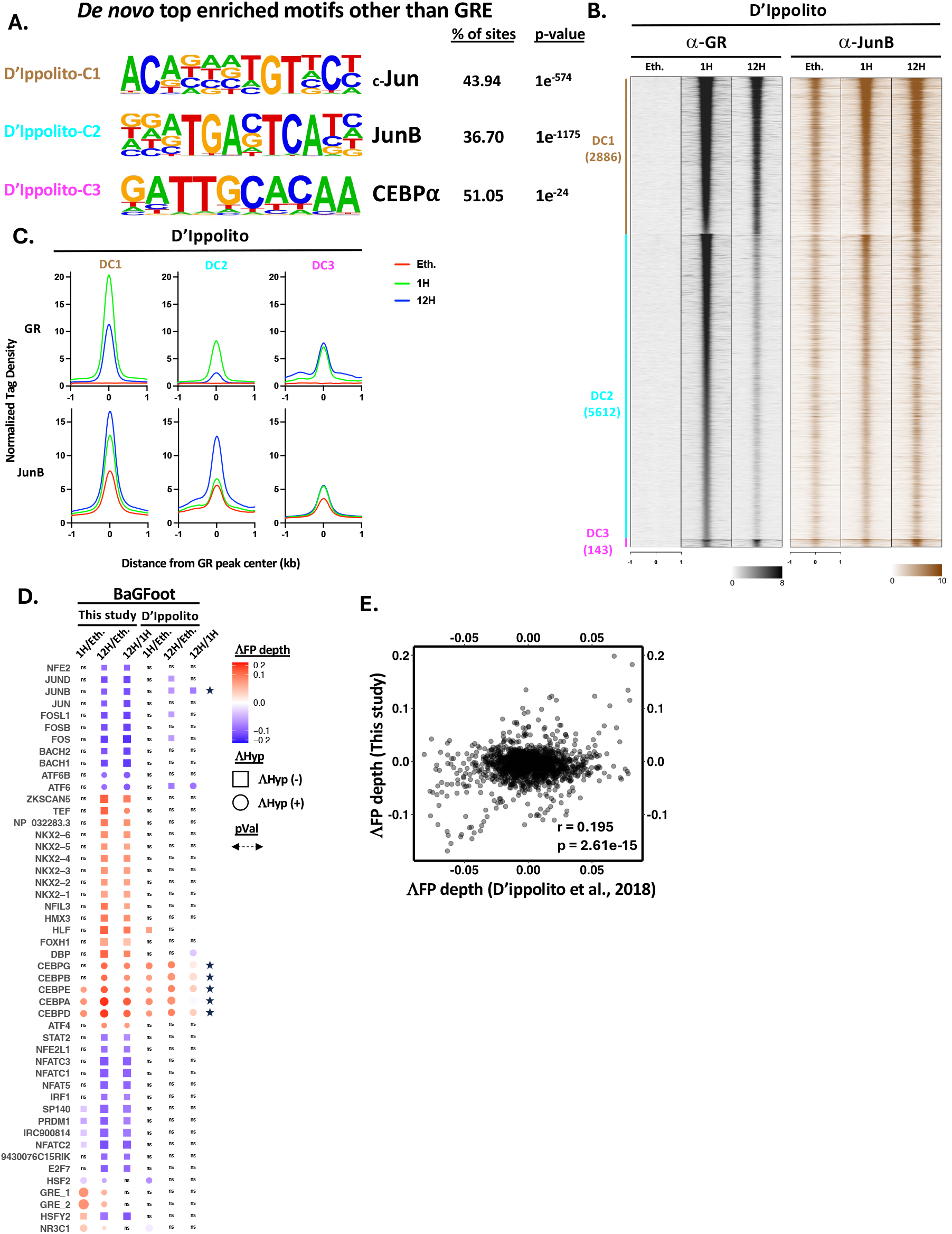
Joint analysis of chromatin accessibility and transcription factor footprinting dynamics by BaGFoot. (**A**) *De novo* motif analysis of GR temporal clusters derived from the D’Ippolito dataset, shown as the percentage of sites containing each motif with associated p-values. (**B**) Heatmaps of GR and JUNB ChIP-seq signals from D’Ippolito *et al*., organized by GR temporal clusters (DC1–DC3) defined using the same peak-calling and clustering strategy applied to the present dataset. (**C**) Aggregate profiles corresponding to panel 5B. (**D**) BaGFoot analysis of the present dataset and the D’Ippolito dataset. Comparisons include 1 h vs Eth., 12 h vs Eth., and 12 h vs 1 h. Heatmaps display changes in footprint depth (ΔFP depth; red to blue), changes in chromatin accessibility (Δ hypersensitivity; squares indicate decreases and circles indicate increases), and statistical significance (symbol width reflects p-value). (**E**) Correlation of footprint depth changes between the present dataset and the D’Ippolito dataset. See also Supplementary Figure 5

Next, we integrated published ChIP-seq data (D’Ippolito et al.) and examined GR and JUNB binding across the temporal clusters defined in the D’Ippolito dataset according to GR dynamic temporal binding (DC1-DC3). Heatmaps and aggregate signal profiles revealed strong GR occupancy at all clusters, with JUNB co-binding prominently observed at early and transient sites (DC1 and DC2), and comparatively reduced signal at late sites (DC3) (Fig. 5B-C). Additionally, JUNB binding dynamics varied across clusters, as evidenced by changes in TagDensity ratios between time points (Supplementary Fig. 5A), further supporting temporally distinct cofactor engagement at GR binding sites. These patterns support a model in which AP-1 factors preferentially associate with early GR binding events.

We next assessed coordinated changes in chromatin accessibility and transcription factor occupancy using BaGFoot^23^. This analysis revealed dynamic shifts in both accessibility (ΔFP accessibility) and footprint depth (ΔFP depth) across comparisons (1 h vs ethanol, 12 h vs ethanol, and 12 h vs 1 h), with distinct transcription factor families exhibiting increases or decreases in occupancy over time (Fig. 5D, Supplementary Fig. 5).

Notably, AP-1-related motifs showed coordinated increase in accessibility and footprint depth at early time points, whereas CEBP family motifs displayed stronger changes at later stages, consistent with their enrichment in the late GR binding cluster (C3). We also analyzed CEBP-β occupancy across the GR temporal clusters defined in the D’Ippolito dataset and observed a progressive increase from the ethanol condition to 12 h of GR activation (100 nM Dexamethasone) in the DC3 (late-binding) cluster (Supplementary Fig. 5B-C).

Importantly, changes in footprint depth were significantly correlated between our dataset and the D’Ippolito dataset (r = 0.195, p = 2.6 × 10^−15^; Fig. 5E), indicating reproducibility and conservation of GR-associated chromatin remodeling dynamics across independent cell lines and species.

### Temporal GR binding is associated with distinct gene expression programs

To determine the functional consequences of GR binding dynamics, we performed temporal transcriptomic profiling following corticosterone treatment. As described previously, multiple dynamic expression profiles were identified among GR-responsive genes^3^. Next, we classified these genes into six distinct expression trajectories based on their temporal response patterns: sustained, transient, or delayed induction, and sustained, transient, or delayed repression (Fig. 6A). Analysis of gene distribution across these categories revealed a predominance of activated over repressed genes, with late-response profiles being the most frequent, followed by transient and then sustained patterns. To rule out a potential confounding effect of transcriptional elongation time, we examined the length of GR target genes and found no significant differences across expression classes (Supplementary Fig. 6A-C).

**Figure 6.**
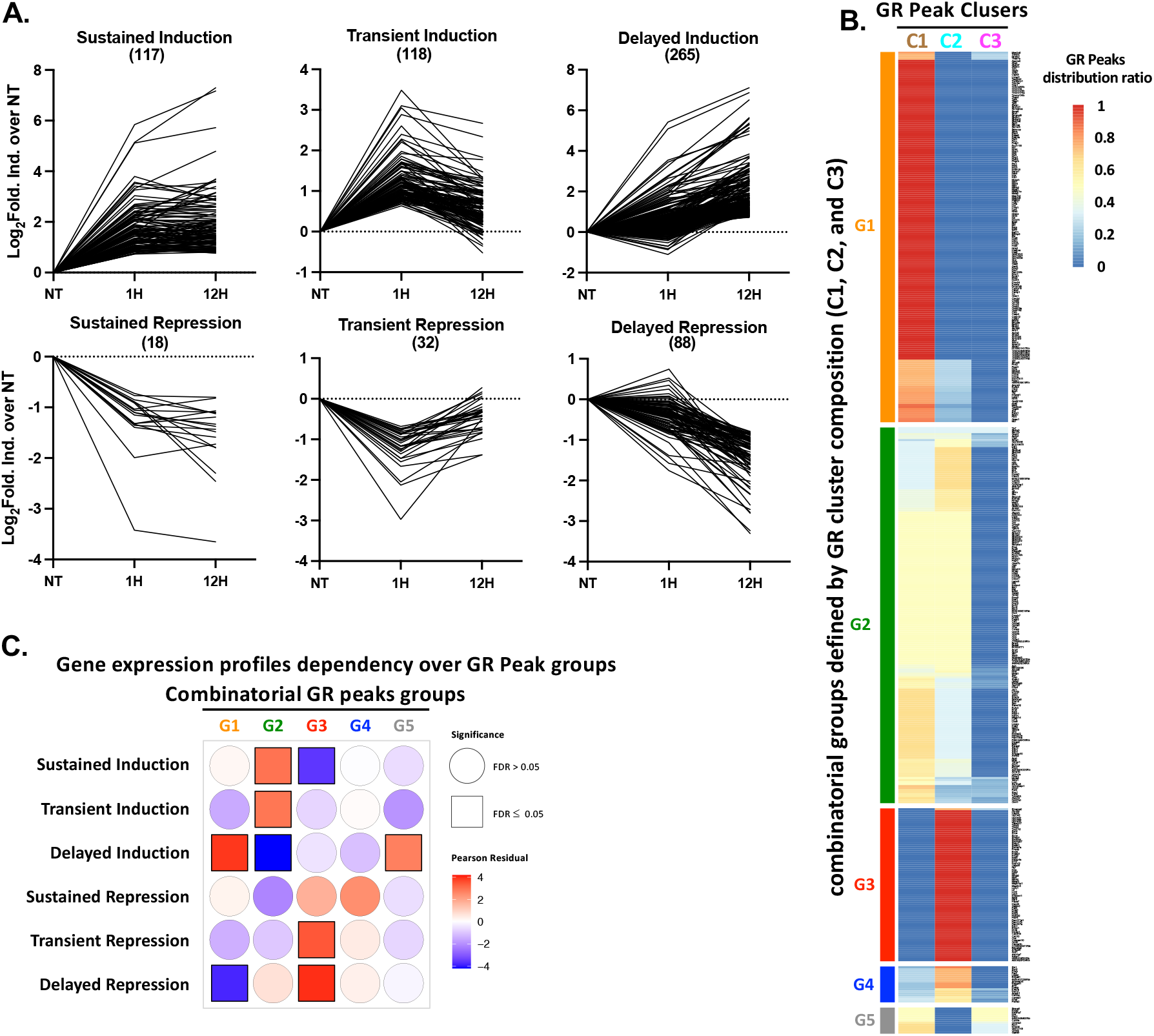
Temporal GR chromatin binding predicts GR-dependent gene regulation profile. (**A**) Temporal GR transcriptome profiling. Data are displayed according to 6 different profiles: induced or activated: sustained, transient or late, based on statistical analysis (p-value < 0.05 and fold change >2). (**B**) Association between GR chromatin binding and gene expression profiles assembled as groups of GR peak clusters (C1-C3). For each gene, GR binding sites were associated thanks to GR HiChIP data^22^ (see method section). (**C**) Pearson correlation analysis between each GR peak cluster groups and gene expression profile (see method section). See also Supplementary Figure 6

We next sought to link these transcriptional programs to GR chromatin binding behavior. Using GR HiChIP interaction data^22^, we assigned each GR peaks to their putative target genes and integrated gene expression data with GR peak clusters (C1-C3) derived from chromatin profiling. While multiple GR peaks can regulate a single gene, we observed that peaks associated with the same gene often originate from different GR binding clusters (C1, C2, and C3). To capture this heterogeneity, we adopted a combinatorial approach in which we quantified the contribution of each GR cluster to individual genes. Specifically, GR peak groups were defined by integrating the relative proportions of peaks from each cluster associated with a given gene, thereby capturing their combinatorial regulatory composition. We then visualized these relationships using a heatmap and assessed gene class enrichment across the resulting combinatorial GR peak groups. Strikingly, the GR peak clusters were specifically enriched in specific gene classes (Fig. 6B). At the level of GR peak groups (G1-G5), clear differences in cluster composition were observed, indicating that combinatorial GR binding patterns contriute to temporal diversity in transcriptional responses.

To systematically quantify these relationships, we performed correlation analysis between GR peak groups and gene expression categories (Fig. 6C). This analysis revealed significant, nonrandom associations between specific GR binding patterns and transcriptional outcomes. In particular, certain GR peak groups were positively associated with induction programs, while others were linked to repression, supporting the idea that temporal and combinatorial GR occupancy encodes distinct regulatory outputs.

Together, integration of GR HiChIP, chromatin profiling, and transcriptomic data reveals that distinct and combinatorial GR binding patterns are nonrandomly linked to specific gene expression programs, giving rise to diverse temporal transcriptional outcomes.

## Discussion

In this study, we define a temporally structured and mechanistically integrated framework for glucocorticoid receptor-mediated gene regulation. Building on prior work demonstrating that GR signaling is dynamic and context-dependent^3,10,11^, we show that GR chromatin binding is organized into conserved temporal classes tightly coupled to chromatin state, cofactor dynamics, and transcriptional outcomes. Together, these findings support a model in which GR signaling is governed by coordinated temporal and combinatorial regulatory mechanisms.

We identify three major temporal classes of GR binding: sustained, transient, and late that are conserved across species and cell types (see Fig. 1). These classes are shaped by intrinsic DNA sequence features and chromatin context. Early and sustained sites are enriched for high-affinity GREs and reside within pre-accessible chromatin, consistent with previous studies demonstrating that accessibility and sequence constrain GR occupancy with high-affinity GREs preferentially located in pre-accessible, nucleosome-depleted regions established by other transcription factors, whereas lower-affinity or nucleosomal GREs require GR engagement with chromatin and subsequent remodeling for stable binding^3,6,30^.

In contrast, late-binding sites show weaker GRE enrichment and lower basal accessibility, indicating a requirement for chromatin remodeling. These results refine chromatin-centric models by showing that the timing of GR binding results from a combination of motif strength and chromatin accessibility, establishing a hierarchical framework for stimulus-responsive occupancy.

Enhancer activation dynamics further distinguish these temporal classes. Among the features examined, H3K27ac most closely tracks GR binding, with rapid enrichment at early sites and delayed accumulation at late sites (Fig. 2A-B). This is consistent with the established role of H3K27ac as a marker of active enhancers deposited by CBP/p300^31,32^ and with prior links between enhancer activity and GR function^9^. The concordance between H3K27ac dynamics and GR occupancy, together with p300 chromatin binding patterns and its transient interaction with GR on the chromatin (Fig. 3B), supports a model in which GR recruits acetyltransferase activity to drive temporally controlled enhancer activation. Notably, chromatin accessibility changes modestly at early-binding sites despite strong GR occupancy (Fig. 2C), consistent with GR engagement at pre-accessible chromatin^6^. In contrast, late-binding sites exhibit coordinated increases in accessibility and acetylation.

In parallel, GR-associated protein interactions undergo substantial temporal remodeling. Proteomic analyses reveal a broad and diverse interactome at early time points that becomes more restricted over time (Fig. 3). This transition aligns with the dynamic nature of GR chromatin interactions observed in live cells^10^ and extends previous chromatin proteomics studies^27^. Together, these results are consistent with a model in which GR-mediated transcription proceeds through distinct phases: an early phase characterized by extensive cofactor recruitment, followed by a maintenance phase with fewer GR-interacting partners. Motif and occupancy analyses further indicate that GR cooperates with distinct transcription factor networks over time (Fig. 4 and 5). Early and transient GR sites are enriched for AP-1 family motifs, including JUNB and ATF3, and display strong JUNB co-binding, consistent with the role of AP-1 factors in priming chromatin for GR recruitment^6,7^. In contrast, late-binding sites are enriched for CEBP motifs and show increased CEBP occupancy, suggesting a distinct regulatory module associated with delayed responses. These findings extend existing models by demonstrating that different transcription factor modules operate at specific temporal phases of GR signaling. Footprinting analyses support this view, revealing coordinated increases in AP-1 occupancy early and CEBP occupancy later. Together, these results indicate that temporally distinct cofactor networks guide GR binding and function.

Notably, integration of GR chromatin binding with transcriptomic profiling via chromatin capture datasets reveals that GR temporal binding patterns are functionally linked to gene expression programs (Fig. 6B-C). Indeed, distinct GR clusters preferentially associate with specific gene sets, and combinatorial GR binding across clusters is nonrandomly linked to defined transcriptional trajectories.

The nonrandom association between GR peak groups and gene expression categories provides quantitative support for a model in which GR functions not as a simple on/oc switch, but as a coordinator of multilayered regulatory inputs. This view is consistent with broader principles of transcriptional regulation, where gene expression outcomes emerge from the integration of binding dynamics, chromatin state, and 3D genome organization^33,34, 30^.

Together, these findings support a unified model in which GR-mediated transcription is governed by the interplay of temporal binding dynamics, chromatin state, combinatorial enhancer activity and specific regulatory complexe assembly. In the early phase, GR binds high-affinity GREs within pre-accessible, AP-1-associated enhancers, driving rapid H3K27ac deposition and broad cofactor recruitment. In the later phase, GR engages weaker, less accessible sites requiring chromatin remodeling and associated with CEBP factors, leading to delayed enhancer activation and reduced protein interactions. Across both phases, combinatorial binding across multiple enhancers specifies gene expression programs, providing a mechanistic basis for temporal diversity in glucocorticoid responses.

The conservation of these temporal patterns across datasets and species underscores the generality of this regulatory architecture and supports the idea that temporal organization of receptor binding and enhancer activation is a fundamental principle of nuclear receptor signaling^6,9^. These findings may have important implications for physiological glucocorticoid responses, in which precise temporal control is essential for processes such as stress adaptation and immune modulation.

Several limitations should be noted. The temporal resolution is limited to two post-stimulation time points, which may not capture intermediate dynamics. HiChIP-based peak-to-gene assignments remain probabilistic, and cofactor associations are largely correlative. Future studies incorporating higher temporal resolution, single-cell approaches, and targeted perturbations of key cofactors such as EP300, AP-1, and CEBP will be essential to establish causality and further define regulatory mechanisms.

In summary, GR-mediated transcription is orchestrated through a temporally ordered and combinatorial framework integrating DNA sequence, chromatin state, cofactor dynamics, and 3D genome organization. This temporal-combinatorial logic provides a mechanistic basis for diverse glucocorticoid responses and establishes a general paradigm for stimulus-responsive gene regulation.

## Acknowledgments

The authors acknowledge the National Cancer Institute Advanced Technology Program Sequencing Facility for sequencing services, as well as the teams at GIGA-ULiège and the NIH High-Performance Computing Systems for their support.

