## Supplementary Figures for "Coordinated Temporal Dynamics of Glucocorticoid Receptor Binding and Chromatin Landscape Drive Transcriptional Regulation"

#### Supplementary Figure 1

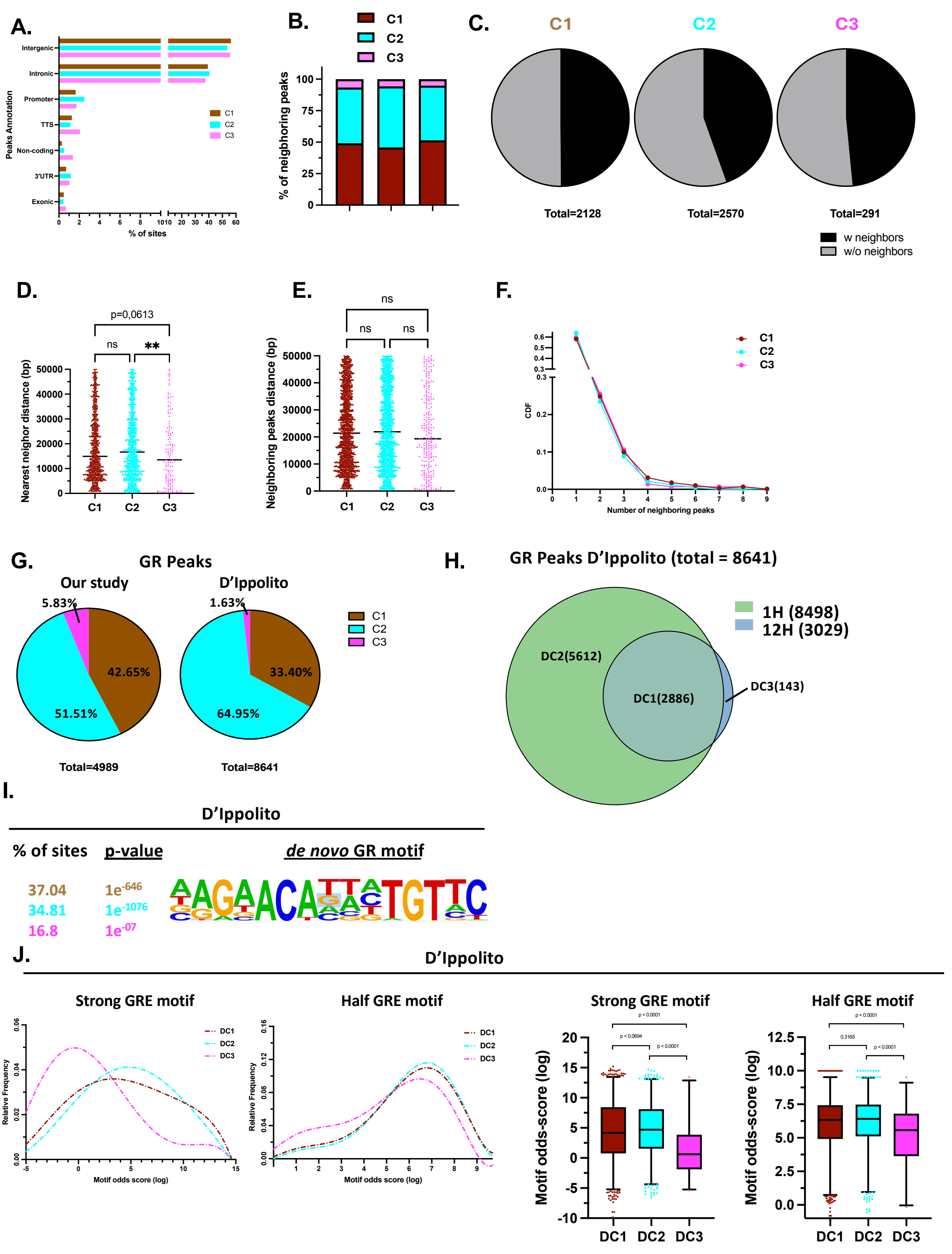


**Supplementary Figure 1. Genomic context and validation of GR peak clusters.** (**A**) Genomic annotation of GR peak clusters from HOMER. (**B**) Cluster identity of GR peaks neighbouring each cluster. (**C**) Fraction of GR peaks with at least one neighbouring GR peak. (**D**) Dot plot of GR peaks nearest-neighbour distances (**E**). Dot plot of GR peaks distances for all neighbouring GR peaks. (**F**) Number of neighbouring GR peaks per peak for each cluster. (**G**). Comparison of cluster distributions between the present dataset and the previously publishes D'Ippolito *et al.* data (D'Ippolito *et al., 2018;* PMID: 30031775). (**H**) Venn diagram of GR peaks detected at 1 h and 12 h dexamethasone in the D'Ippolito dataset. GR peak clusters are names DC1, DC2 and DC3 analogously to Fig. 1D. (**I**) Top enrichment GR motif from a *de novo* motif analysis of all GR peaks from D'Ippolito et al (n = 8641). The enrichment is displayed as percentage of the number of sites with the calculated p-values. (**J**) Analysis of canonical (strong) and half-site (weak) GR response elements motifs odds scores (log scale) and frequencies were calculated for each GR cluster. Boxplots of motif odds-score distributions from panel G (KW-test) p-val<0.05 is considered significant.

### Supplementary Figure 2

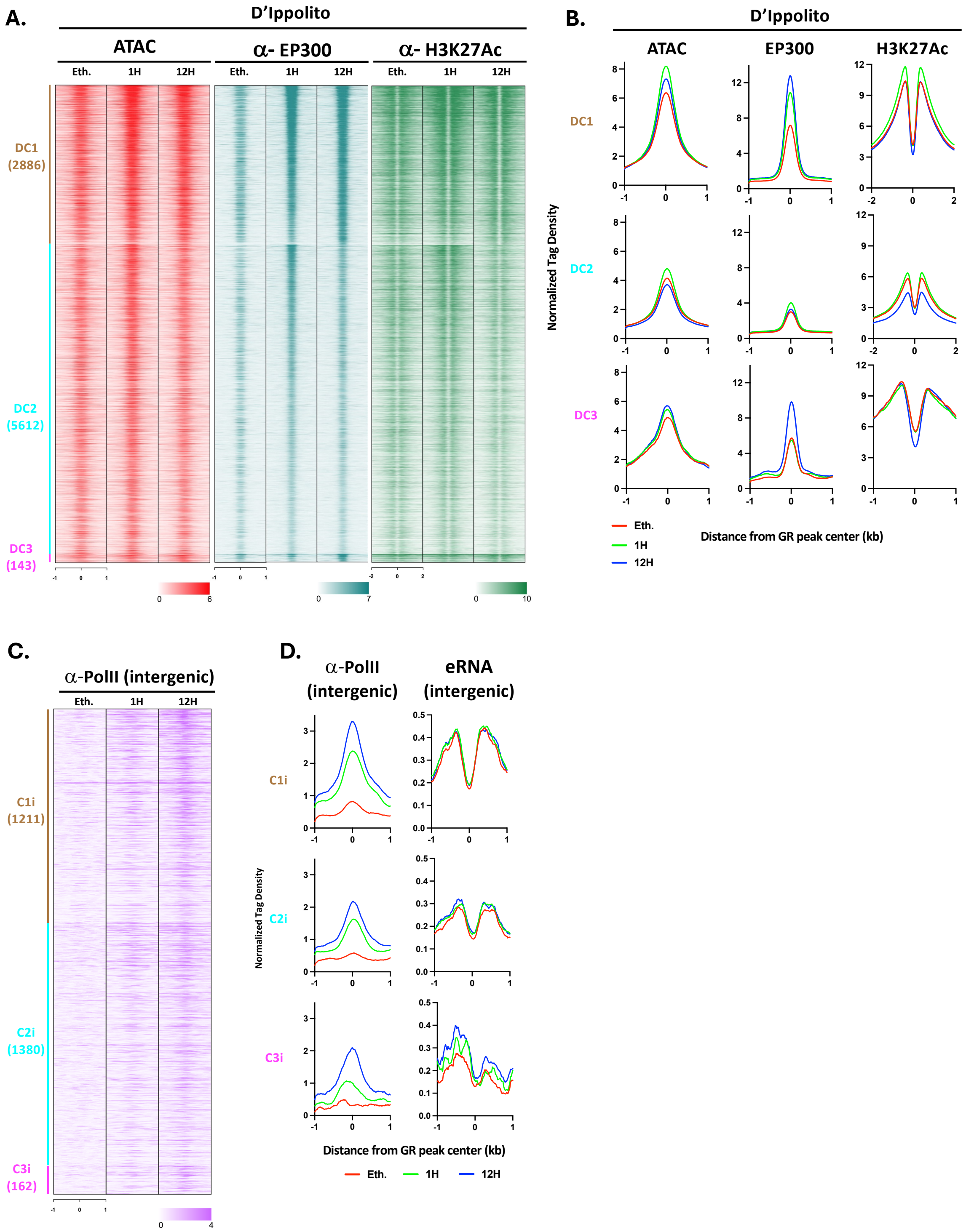


**Supplementary Figure 2. RNA polymerase II occupancy at intergenic GR binding sites.** (**A**) Heatmaps of ATAC-seq, EP300, and H3K27ac Chip-seq signals at GR binding sites, shown ±1 or 2 kb from peaks center and organized by GR temporal clusters (DC1-DC3). (**B**) Aggregate plots related to panel S2A for each cluster profiles (20-bp bins) around peaks center. (**C**) Heatmaps of Pol II ChIP-seq signal at intergenic GR binding sites, displayed ±1 kb from peaks center and grouped by intergenic cluster (iC1–iC3). (**D**) Aggregate profiles corresponding to panel S2C, calculated as summed read density in 20-bp bins within ±1 kb of peaks center.

### Supplementary Figure 5

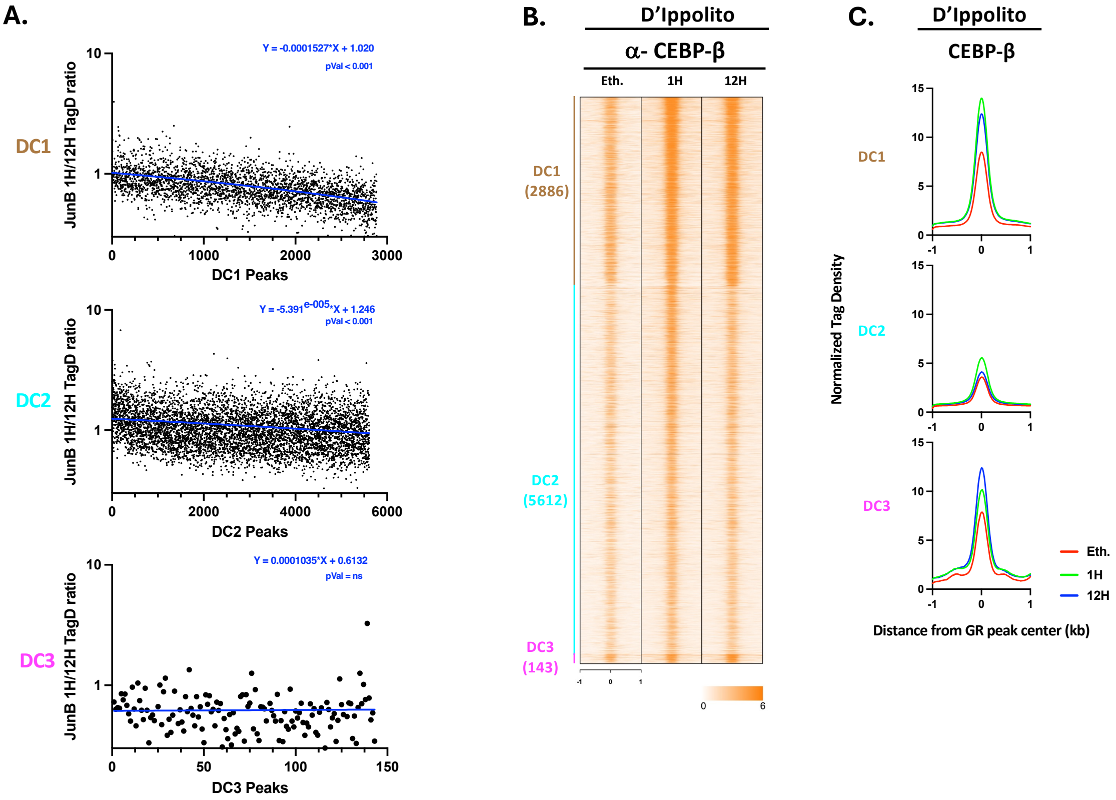


**Supplementary Figure 5. Chromatin landscape of the GR temporal peak clusters in the D'Ippolito dataset.** (**A**) JunB TagDentisty ratio ordered 1 h vs 12 h displayed for each GR cluster ordered by GR peak score. (**B**) Heatmaps of CEBP-β Chip-seq signals at GR binding sites, is shown ±1 or 2 kb from peaks center and organized by GR temporal clusters (DC1-DC3). (**C**) Aggregate plots related to panel S5B for each cluster profiles (20-bp bins) around peaks center.

### Supplementary Figure 6

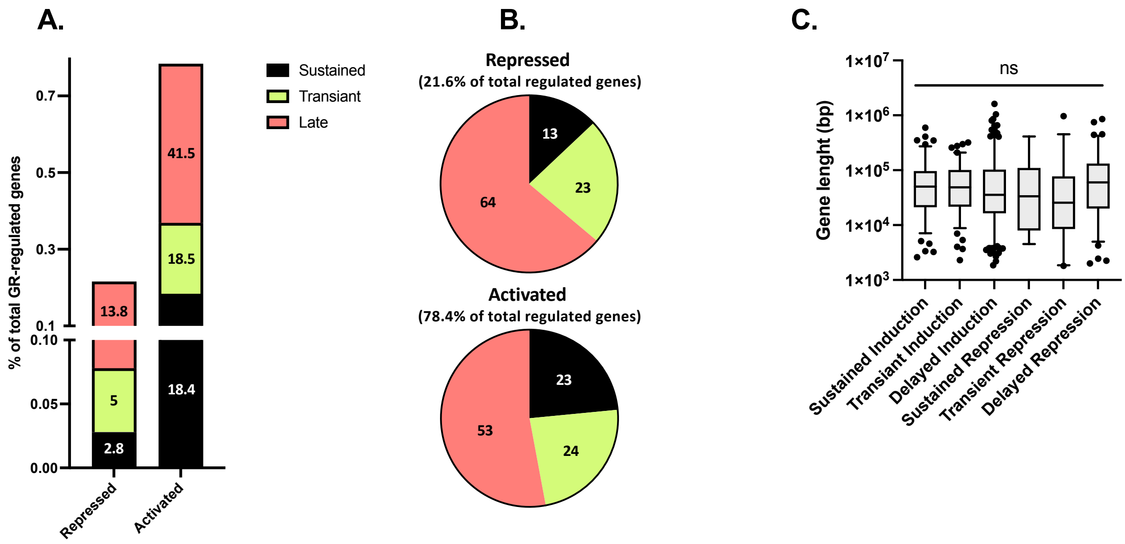


**Supplementary Figure 6. Gene expression profile investigations** (**A**) Proportion of each gene profiles as compared to the overall number of analysed genes. (**B**) Venn diagram related to S6A (**C**) Gene length analysis for each gene profiles.
